# Pulsed electrical stimulation enhances intestinal permeability

**DOI:** 10.64898/2026.06.09.731152

**Authors:** Batoul Khlaifat, Heba Naser, Erika Sargasyan, Abdel-Hameed Dabbour, Salma Nassar, Khalil B Ramadi

## Abstract

Oral ingestion of drugs remains the most convenient method for pharmacotherapy. However, oral absorption is hampered by digestive enzymes and the intestinal epithelial barrier. Here we investigate the ability of electrical stimuli to biologically modulate intestinal permeability. We demonstrate that pulsed electrical stimulation increases intestinal permeability, facilitating transport of chemical species across the epithelium. We evaluate the effects of several stimulation parameters *in silico* and subsequently characterize the biological effects in vitro using Caco-2 colorectal cancer cells, and an acute in vivo intestinal model. Our findings suggest that these effects may be mediated through calcium-dependent interactions with tight junction proteins which induce a reversible permeability increase differing based on the total charge delivered, amplitude and frequency of the current delivered. Pulsed electrical stimulation could be a potential strategy for transiently modulating the intestinal barrier.

## INTRODUCTION

The universally preferred method of drug administration is the oral route. Oral pharmacotherapies are absorbed through the intestinal epithelia and then circulate systemically. Epithelial uptake is enhanced through intestinal villi and microvilli, which increase surface area for absorption [1]. Oral drug delivery, however, remains limited in many cases due to the selective permeability of the epithelial barrier and many drugs are not readily absorbed.

The barrier function of the intestinal epithelia protects against luminal pathogens and unwanted chemicals, but also restricts drug transport. Epithelial tight junctions primarily control paracellular transport [2]. Chemical permeation enhancers such as surfactants, cyclodextrins, and nanoparticles can be co-delivered with drugs to facilitate paracellular transport, although issues scalability, biocompatibility, and complexity persist [3, 4].

Electrical manipulation of tissues has been shown to regulate proliferation, differentiation, and cell-cell communication [5-8]. Ingestible electroceuticals can deliver electrical stimuli throughout the GI tract for modulation of motility and metabolism. The use of electrical stimuli to facilitate drug delivery has largely been in the form of iontophoresis, where an applied electric field electrophoretically drives charged molecules through tissue. In transdermal and commercially available implementations the two electrodes are typically positioned on the same side of the barrier, with current passed through the tissue between them, rather than across electrodes placed on opposing faces of the membrane [9-11]. However, this is challenging to implement in the gut as ingestible devices do not continuously contact the epithelium, and when they do so only contact the apical side[12]. One of the few studies which propose iontophoretic-based stimulation through the luminal intestinal space utilizes hydrogel adhesive patches that stick to the intestinal wall, and deliver drugs through a reservoir that iontophoretically drives drug transport [13].

Here, we show how applying non-uniform, pulsed electrical current to the apical epithelium can transiently and reversibly modulate intestinal permeability. We apply this in vitro in a Caco-2 cell model and find that the extent of barrier permeability changes correlates with amplitude, frequency, and overall stimulus duration. We utilize an acute in vivo model externalizing the intestine and find a similar increase in permeability following electrical stimulation. Electrical pulses accomplish this by interfering with calcium signaling to induce changes in tight junction proteins. Focal, transient delivery of electrical stimuli could be one approach to enhance oral drug absorption.

## RESULTS

### Computer simulations

We developed COMSOL Multiphysics simulations to understand the electric fields generated by iontophoretic stimulation (IPS) and direct stimulation (DS) as we varied parameters of the current applied. For both, we modeled 2-dimensional plane of epithelial cells grown on a transwell in vitro. Direct stimulation utilized two platinum (Pt) electrodes spaced 3.6 mm apart above the apical cell layer. Iontophoretic stimulation electrodes were placed such that one electrode is above the cell layer and the ground is modeled below the cell layer and transwell (Figure 2A-F). 2-dimensional model geometries were modeled after the standard 0.3 cm^2^ transwell dimensions (Figure 2B). In both setups, electrodes were not directly in contact with the cell layer.

**Figure 1.**
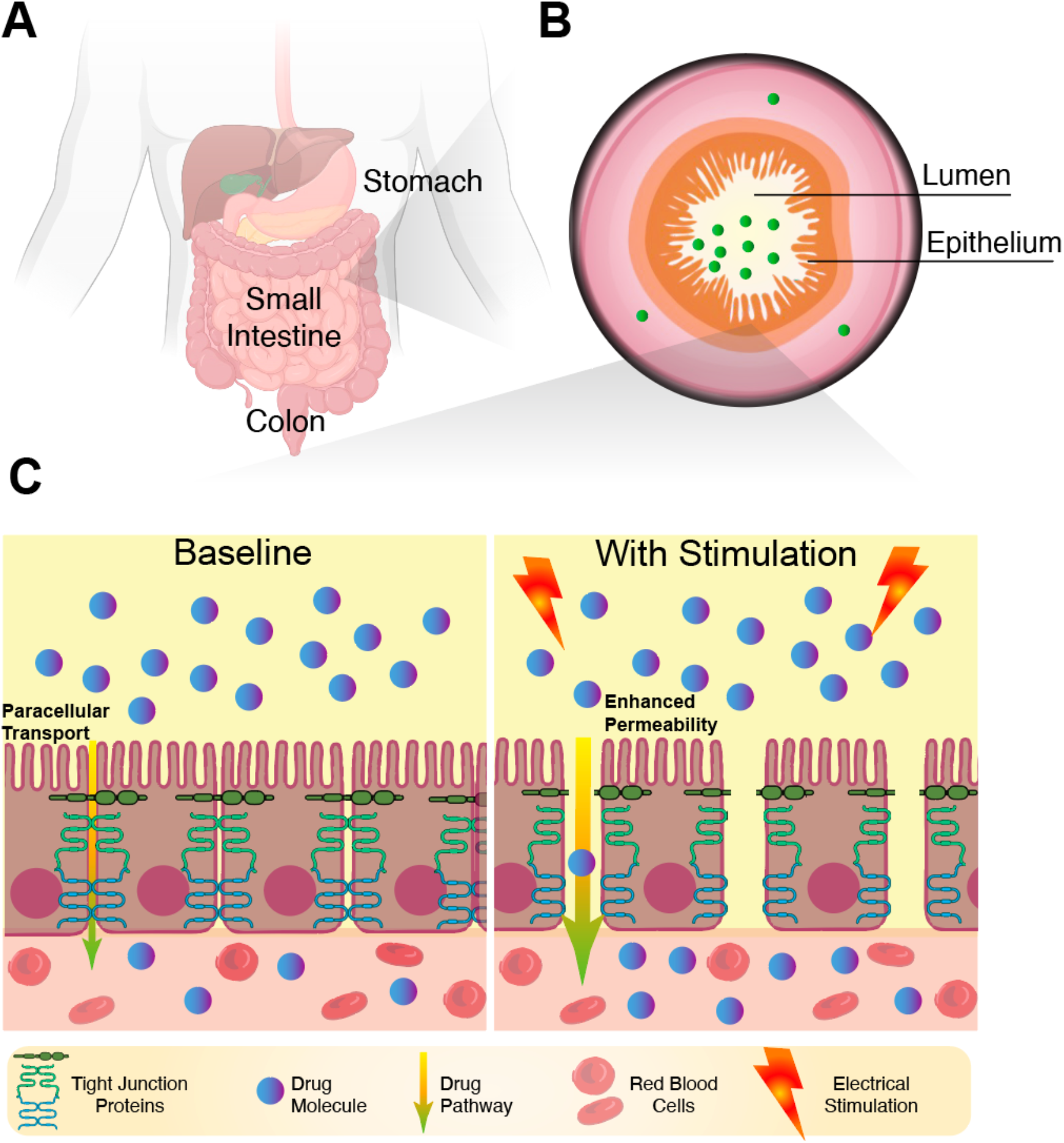
(A) Schematic of human gastrointestinal tract, (B) Schematic of intestinal cross-section, (C) Epithelial cells and their associated tight junction proteins responsible for paracellular transport at baseline and following electrical stimulation showing enhanced permeability.

**Figure 2.**
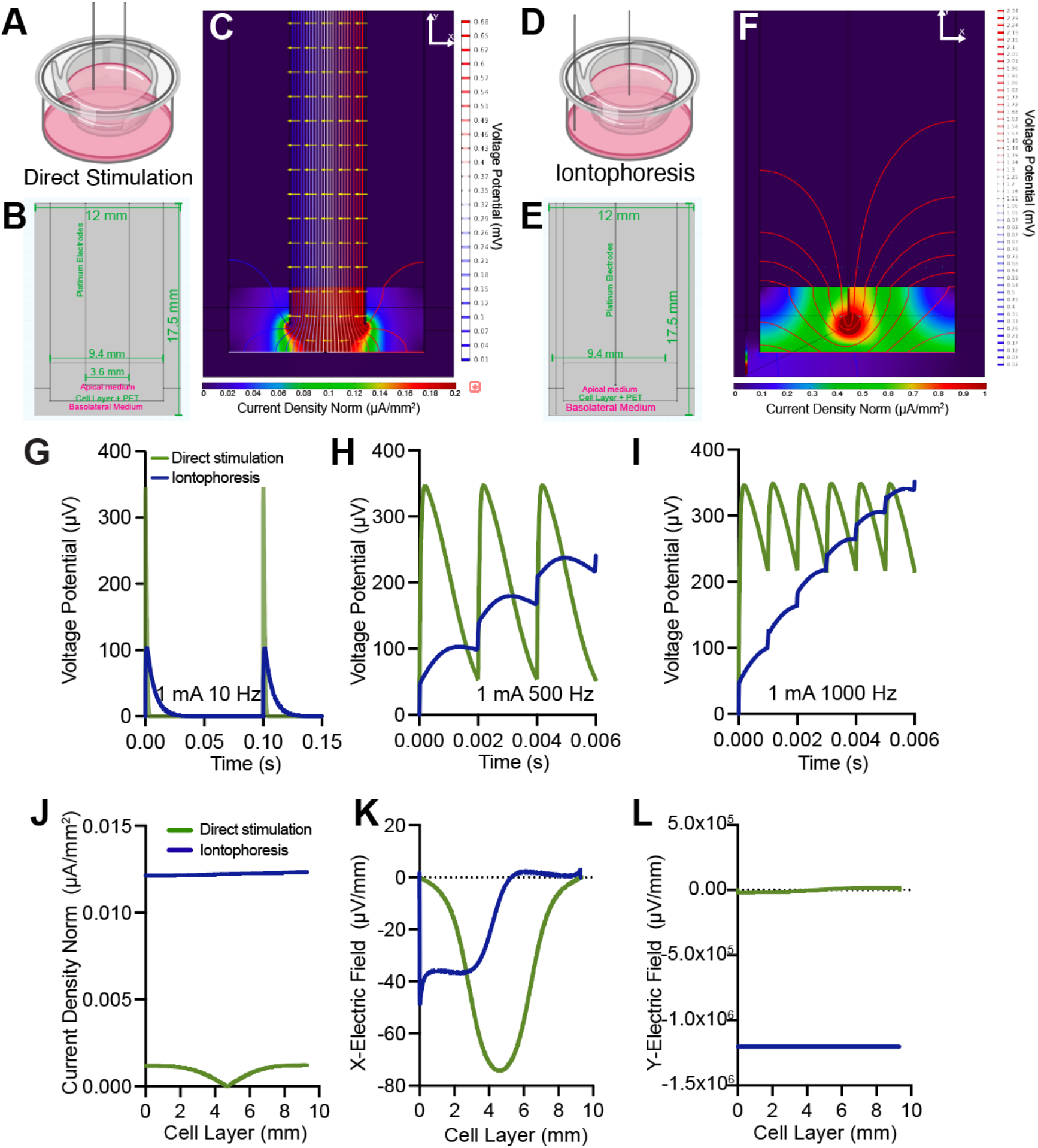
(A) Schematic of in vitro Direct Stimulation setup, (B) Geometric View and dimensions of Direct Stimulation Model, (C) 2D COMSOL Simulation of the in vitro direct stimulation, (D) Schematic of in vitro Iontophoretic stimulation, (E) Geometric view and dimension of Iontophoresis model, (F) 2D COMSOL Simulation of the in vitro iontophoretic stimulation, (G-I) Voltage potential in stimulated cell layer vs Time at a frequency of (G)10 Hz, (H) 500 Hz, and (I) 1000 Hz. (J) Current Density Norm of bulk cell layer at 1 mA 500 Hz stimulation, (K) X-Component Electric field in the bulk cell layer at 1 mA 500 Hz stimulation (L) Y-Component Electric field in the bulk cell layer at 1 mA 500 Hz stimulation

We applied pulsed electrical current (500 Hz, pulse width 1 ms) in both setups, varying the frequency and amplitude to investigate its effects on bulk cell layer model (Figure S1). Voltage potential concentrates more between the electrodes in direct stimulation, while current density (surface plot) is higher in the medium during iontophoretic stimulation (Figure 2C,2F). Increasing amplitude from 10 µA to 5 mA increases voltage potential at the cell layer (Figure S2). We probed the cell layer voltage potential as a function of time in 1 mA applied current while varying the frequency (10 Hz, 500 Hz, 1000 Hz) (Figure 2G-I).

Under 10 Hz stimulation, direct stimulation produces a larger peak voltage amplitude compared with iontophoresis. While both voltage profiles decay to baseline between pulses, direct stimulation produces more rapid decays than iontophoresis (τ = 0.001025 vs 0.006550, respectively) (Fig. 2G). This difference is amplified as with increased stimulation frequency. Increasing pulse frequency to 500 Hz leads to an accumulation of voltage in IPS, characterized by a stepwise increase in baseline voltage across stimulation cycles (Figure 2H). Direct stimulation, however, presents a consistent gradual decay to a baseline of ≈ 50 µV and a higher pulse amplitude than the initial amplitude of IPS pulses. The divergence between stimulation modalities becomes more pronounced at 1000 Hz frequency (Figure 2I). In IPS, peak voltage of the cell layer rises continuously while direct stimulation pulses peak at consistent levels throughout cycles. However, even in DS the reduced inter-pulse interval limits complete discharge between cycles.

The iontophoretic potential approaches a quasi-steady elevated voltage state, suggesting that the system behaves increasingly as a charge-accumulating interface at higher frequencies. These results are consistent with the known resistive-capacitive behavior of biological systems at higher stimulation frequencies. The ability of the modeled cell layer to discharge between pulses is strongly dependent on stimulation frequency and differed substantially between stimulation configurations. The IPS model exhibited greater apparent resistance, resulting in slower discharge kinetics and progressive charge accumulation across repeated pulses. Direct stimulation maintained a predominantly transient and pulse-following electrical response, marked by its pulse recovery.

We next mapped the spatial distribution of the electric field and the current density magnitude in the IPS and DS configurations during a stimulation paradigm of 1 mA, 500 Hz. The current density at the cell layer, is reduced in DS (≈ 0.001 µA/mm^2^) compared to IPS (≈ 0.01 µA/mm^2^) (Figure 2J). The X-electric field at the cell layer during direct stimulation localizes to the region under the electrodes reaching a maximum value of ≈ -73 µV/mm, whereas the electric field in IPS exhibits lower localization at the periphery of the cell layer, in line with the trajectory between the working electrode and the basal electrode (Figure 2K). The Y-electric field in DS reveals minimal contribution compared to IPS where the field averages to -1×10^6^ µV/mm (Figure 2I). These results highlight the disproportionate effect of the vertical and horizontal components in both stimulation paradigms, and emphasizes the predominance of the X horizontal component of the direct stimulation configuration, relative to the iontophoretic model which relies primarily on the high values of the vertical component.

### Electrical characterization of caco-2 cells

We constructed the electrical delivery apparatus as a series circuit with a pulsed stimulation source assuming the equal distribution of current throughout the wells (Figure 3A). Biological receiver was represented by an equivalent resistive-capacitive (RC) model, in which the Transwell membrane was modeled as a resistor while the Caco-2 cell monolayer and the culture medium as capacitive elements, collectively governing the impedance and temporal distribution of the applied current. Electrical stimulation was delivered through two Pt-Ir electrodes (0.004”) spaced **0.3 cm** apart and submerged in cell medium (Figure 3B-C).

**Figure 3.**
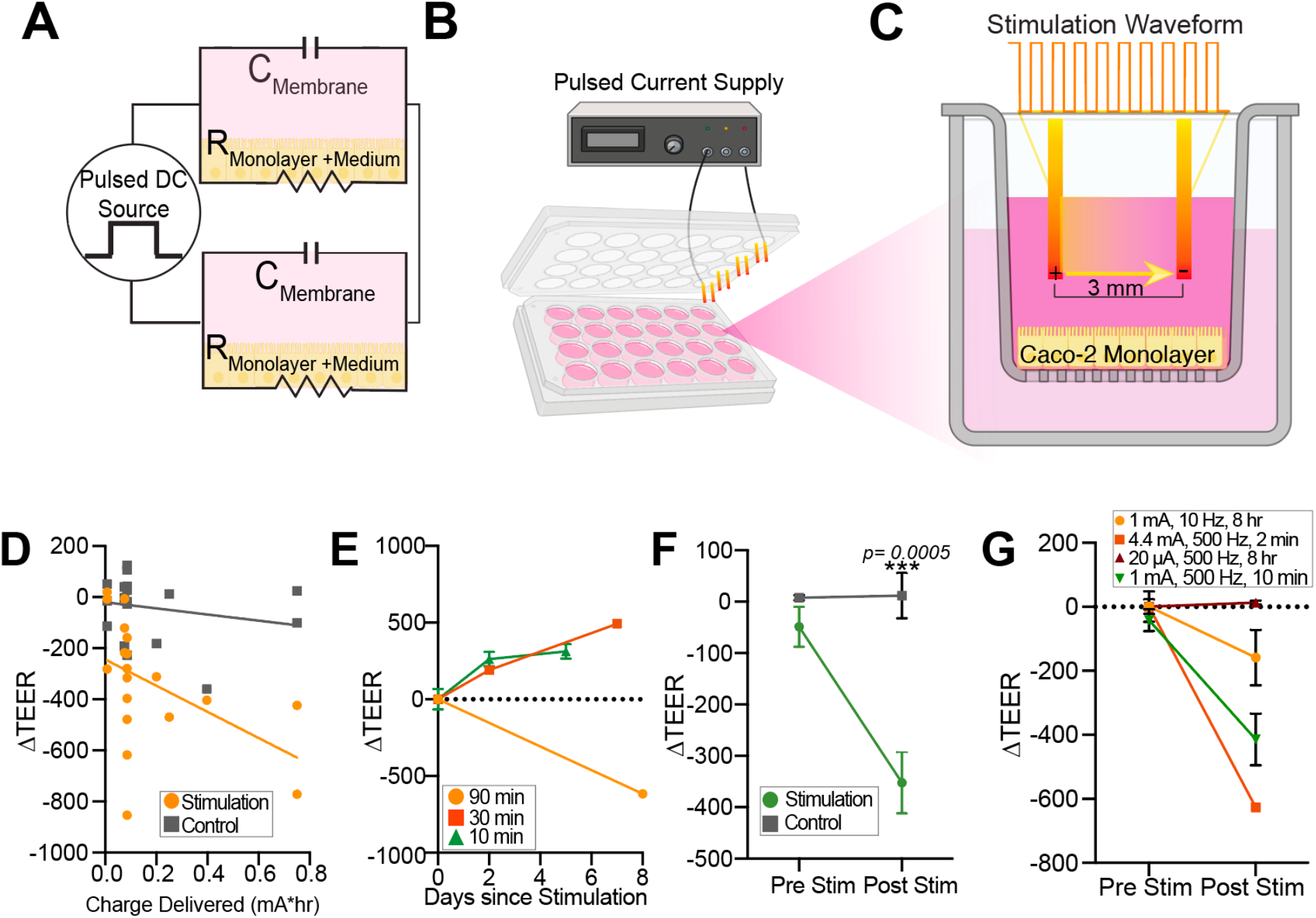
(A) Circuit Diagram of Electrical Stimulation Set Up, (B) Schematic of Electrical Stimulation Lid used during In vitro stimulation of Caco-2 cells, (C) Schematic of Caco-2 intestinal model grown on Transwell insert with Platinum electrodes and delivered square waveform, (D) Change in transepithelial electrical resistance (TEER) of Caco-2 cells at various delivery of electric charges, (E) Recovery of TEER values after stimulation at different electrical parameters, (F) change in TEER after 1 mA, 500 Hz and 10 mins of electrical stimulation, (G) change in TEER after variation of electrical stimulation parameters whilst maintaining charge delivery to Caco-2 cells.

To understand the effects of pulsed electrical current stimulation on the Caco-2 monolayer’s permeability, we electrically stimulated cells at various parameters and compared the change in transepithelial electrical resistance with the overall charge delivered (mA* hr), consolidating the different stimulation conditions into a single quantitative metric (Table 1). A decrease of 800 Ω in TEER displayed irreversible cell detachment and damage and was therefore defined as the threshold which was measured when a 0.8 mA* hr was delivered to the cells. Overall, the extent of TEER decreases correlated with the total charge delivered (Figure 3D).

**Table 1:** Total charges delivered and the parameters (amplitude, frequency, pulse width, duration) used in the electrical stimulation of caco-2 cells.

| Charge Delivered ($\text{mA} \cdot \text{hr}$) | Parameters (*Amplitude, frequency, pulse width, overall stimulation duration*) |
| --- | --- |
| $7.5 \times 10^{-3}$ | 10 $\mu\text{A}$, 500 Hz, 1 ms, 90 min |
| $7.5 \times 10^{-2}$ | 100 $\mu\text{A}$, 500 Hz, 1 ms, 90 min |
| $8.5 \times 10^{-2}$ | 1 mA, 500 Hz, 1 ms, 10 min |
| $2 \times 10^{-1}$ | 1 mA, 100 Hz, 1 ms, 120 min |
| $2.5 \times 10^{-1}$ | 1 mA, 500 Hz, 1 ms, 30 min |
| $3.97 \times 10^{-1}$ | 500 $\mu\text{A}$, 500 Hz, 1 ms, 90 min |
| $7.5 \times 10^{-1}$ | 1 mA, 500 Hz, 1 ms, 90 min |

Next, we stratified three different durations of the same parameters to determine whether different electrical stimulation charges caused lasting impairment to monolayer integrity. Caco-2 monolayers were stimulated using identical parameters (1 mA, 500 Hz, 1 ms pulse width) for 90, 30 or 10 minutes (Figure 3E). The 90 minute-stimulation resulted in a sustained reduction in TEER that persisted for up to 8 days post-stimulation, with no evidence of recovery. In contrast, monolayers stimulated for 30 minutes or 10 minutes exhibited a gradual restoration of TEER over a comparable time frame. As the 10-minute stimulation achieved barrier modulation while allowing subsequent recovery, we selected this as the stimulation paradigm for further studies. 10min stimulation decrease transepithelial resistance by 400 Ω • cm^2^ (Figure 3F).

We then probed whether varying the parameters while maintaining total charge delivered (0.08 mA*hr) would induce similar changes to TEER. We maintained the same pulse amplitude (1 mA) and decreased frequency to 10 Hz for a total of 8 hours stimulation. We then increased the current amplitude (4.4 mA) and decreased total stimulation time to 2 mins at the same 500 Hz frequency. We also tested a lower current amplitude stimulation (20 µA) over 8 hours while maintaining frequency of 500 Hz. This set up allows us to examine contributing effects of each stimulation parameter to TEER.

Decreasing stimulation frequency to 10 Hz (over 8 hr) resulted in a smaller change to TEER, compared to 500 Hz stimulation (180 vs 350 Ω•cm^2^). Increasing amplitude over a shorter time, on the other hand, resulted in a TEER change of ∼700 Ω•cm^2^. Decreasing 20 µA amplitude at the original (500 Hz) frequency and 8 hours duration did not induce any significant change in the cell monolayer’s electrical resistance (Figure 3G).

This suggests that TEER change is not uniquely dependent on the total charge delivered during the stimulation window. Biological response is more sensitive to current amplitude than frequency or overall duration of stimulus. In line with the computational results (Figure 2G-I), lower frequency (10 Hz) also allows the system sufficient relaxation time to return to a baseline voltage between pulses and could contribute to the reduced change in TEER observed experimentally. Prolonged stimulation at lower amplitude and frequency has the least effect on the TEER, suggesting that tight junction disruption also has a minimum amplitude threshold.

### Biological characterization of stimulated caco-2 monolayers

We next investigated the effects of electrical stimulation on the chemical permeability of the monolayer. We applied 10 mins of 1 mA 500 Hz stimulus to caco-2 monolayers and measured the rate of fluorescein transport across the monolayer for 120 minutes (Figure 4A). We then normalized the apparent permeability against controls. The electrically stimulated monolayer was 2x more permeable than the unstimulated controls (Normalized P_app_ ≈ 2.11, (Figure 4B)). These findings are consistent with the previously observed decrease in TEER, confirming that electrical stimulation compromises barrier integrity in a biologically meaningful manner.

**Figure 4.**
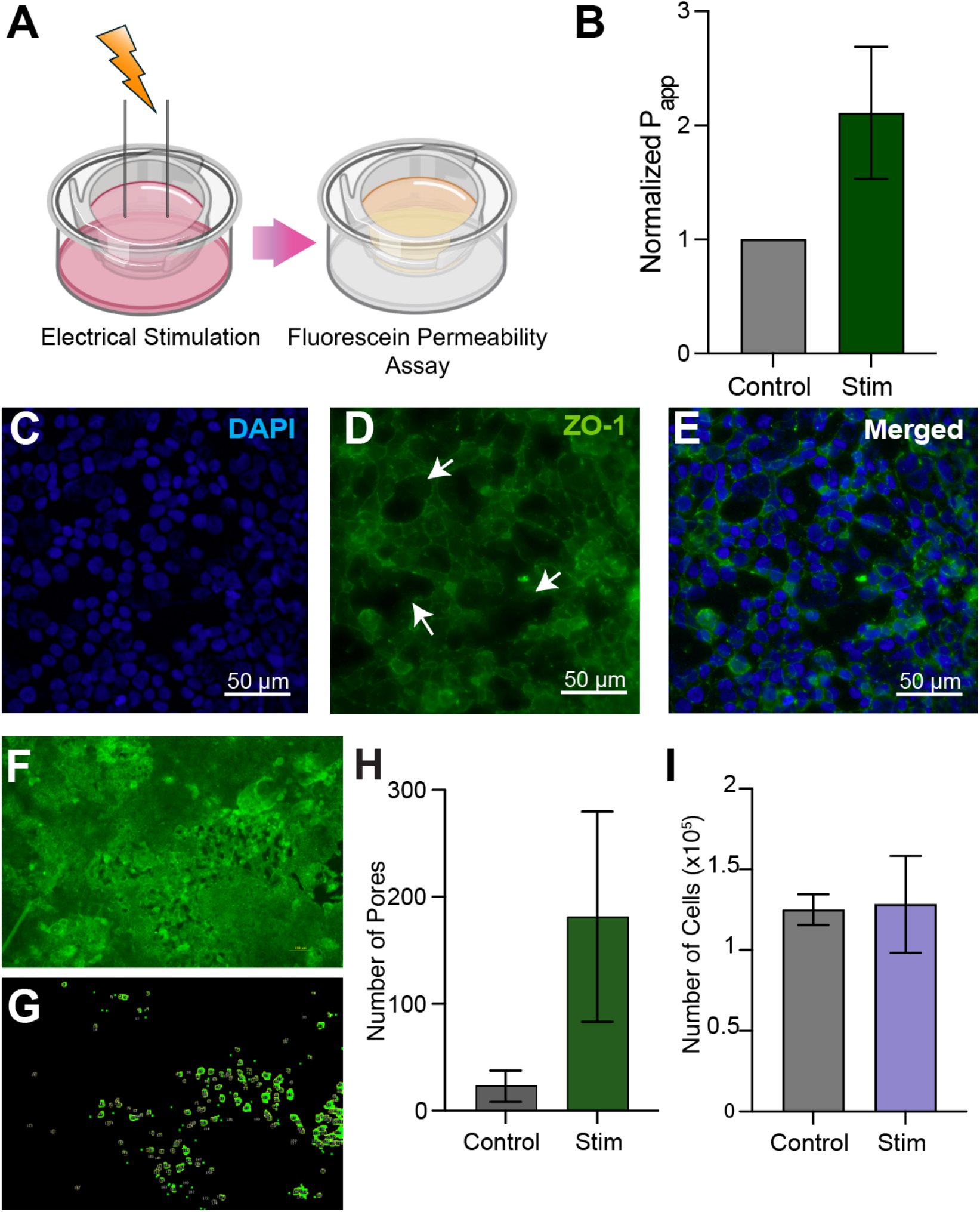
(A) Schematic of permeability assay, (B) Normalized Apparent Permeability of fluorescein in control sample and stimulated sample of Caco-2 cells, (C,D) (C) DAPI and (D) ZO-1 stained electrically stimulated caco-2 cells, (F) Electrically stimulated Caco-2 cells pores, (G) Representative image analysis of electrical stimulation-induced pores in cells, (H) Pore count in control vs. stimulated cells, (I) Cell count of control and stimulated cells.

We then examined the morphological changes that arise in tight junction proteins after stimulating the monolayer. We stained for zonula occludens 1 protein and stained DAPI to investigate cell death (Figure 4C-E). Visible pores were quantified and counted against the control monolayers using FIJI/ImageJ (Figure 4F-G). Stimulated Caco-2 monolayer showed an 8-fold increase in the pore count across the analyzed monolayers, with no difference in overall cell count (Figure 4H,I). While electrical stimulation seems to impact tight junction morphology in caco-2 monolayers, this does not appear to induce cell death.

### Calcium role in intestinal permeability modulation

We sought to probe the mechanism through which electrical stimulation modulates tight junctions, and we examined the role of calcium ions in this process. We hypothesized that an increase in cytoplasmic calcium driven by the flux of extracellular calcium and consequent release of calcium ions from the endoplasmic reticulum promotes phosphorylation of tight junction proteins such as ZO-1 thereby altering the paracellular permeability of the intestinal model (Figure 5A).

**Figure 5.**
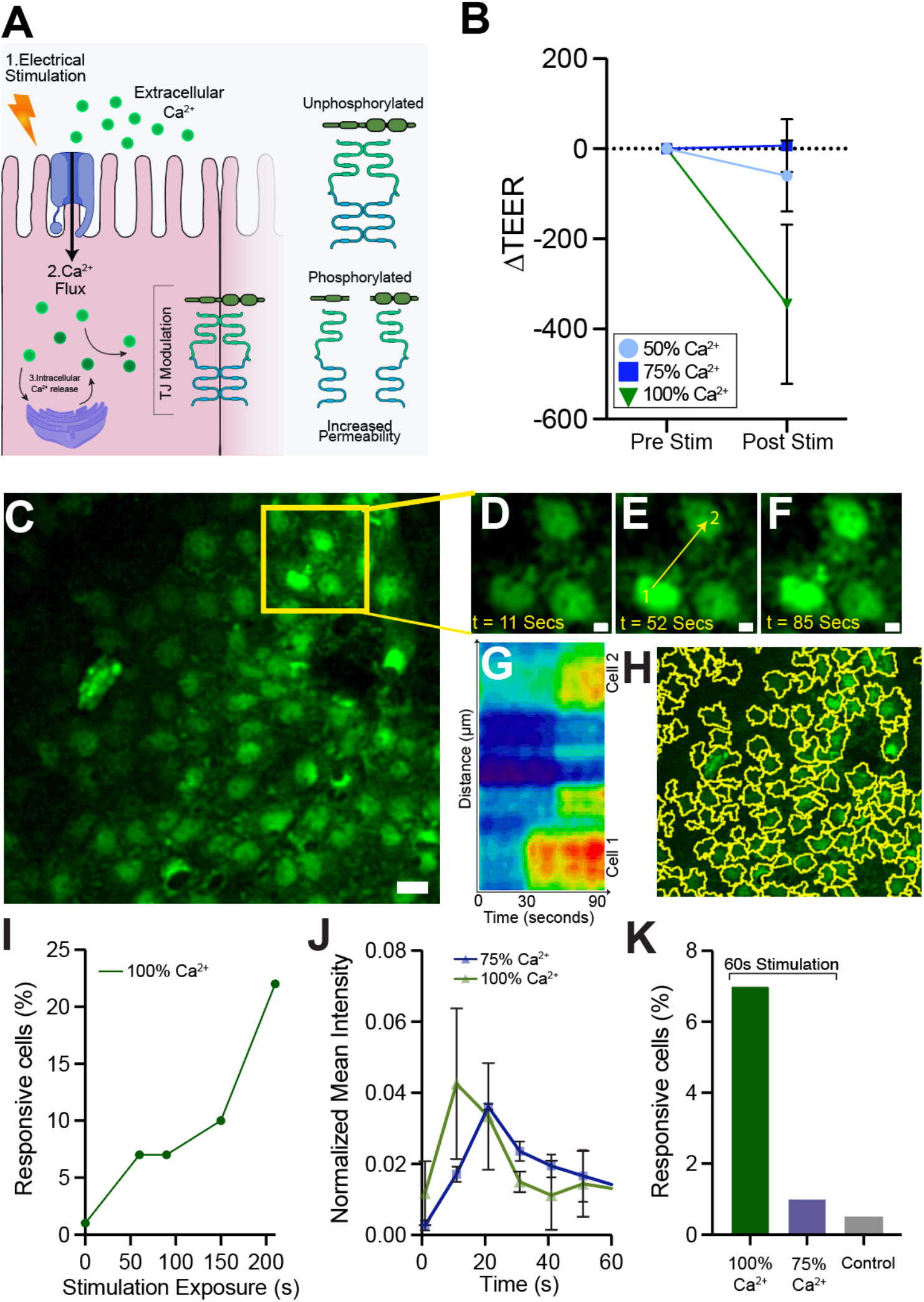
(A) Schematic of Calcium Flux and interaction with Tight Junction Proteins, (B) Change in Transepithelial electrical resistance of Caco-2 cells with 50%, 75%, 100% Calcium medium, (C) Live Calcium Imaging of Caco-2 cells (Scale bar = 20 µm), (D-F) Caco-2 Cells imaged (D) before, (E) during, and (F) after electrical stimulation (Scale bar = 5 µm). (G) Kymograph of cell signaling of cells in D-F (1 px = 0.3 µm;1.3 s), (H) Cellular segmentation using ImageJ. (I) Percentage of responsive cells in cell populations with stimulation exposure in 100% Calcium, (J) Normalized Mean intensity of aligned peaks in cells electrically stimulated in 75% and 100% calcium medium. (K) Percentage of responsive cells in 60 seconds stimulated cells in 100% calcium and 75% calcium and non stimulated controls.

To assess how partial calcium depletion influences the response of electrically stimulated Caco-2 cells, we applied stimulation (1 mA, 500 Hz,10 minutes) to Caco-2 cells in 75% and 50% calcium HBSS medium and compared the change in TEER to the standard 100% calcium medium. In cells stimulated under 100% calcium conditions, the TEER decreased by ≈ 200 Ω/cm^2^. In contrast, the TEER of stimulated cells in 75% calcium medium remained near baseline (6.675 Ω/cm^2^, Dunnet-adjusted p= 0.0206*). Notably, stimulation in 50% calcium resulted in a moderate decrease in TEER (≈ -60 Ω/cm^2^, Dunnet-adjusted p = 0.0593 ns), suggesting that increased depletion of extracellular calcium could also impact cellular permeability (Figure 5B).

To further investigate calcium signaling dynamics in electrically stimulated caco-2 cells, we performed live cell imaging of Caco-2 cells during electrical stimulation (1mA 500 Hz) applied in intervals (30-60 seconds) (Figure 5C). Figure 5D-F shows a heterogenous and localized calcium oscillation induced by stimulation when applied at T= 30 seconds. The kymograph in Figure 5G illustrates calcium signal propagation between these two neighboring cells during electrical stimulation, with cell 2 activating shortly after the signal intensifies in cell 1. Spontaneous oscillations are also exhibited in certain population, further illuminating the heterogenous response during electrical stimulation (Figure S3).

We segmented the imaged populations, yielding an average of ≈ 338.2 cells per imaging field (Figure 5H), and filtered normalized calcium signals based on responsiveness (SNR ≥ 0.1). Repeated stimulation in 100% Calcium medium increased the percentage of responsive cells, reaching 25% after 200 seconds of cumulative exposure (Figure 5I).

Next, we compared calcium dynamics in 100% calcium vs 75% calcium medium. The peak-aligned traces show similar mean intensities (≈ 0.04 au) for cells stimulated in both groups during 30 seconds of stimulation (Figure 5J). However, when comparing the percentage of responsive cells in populations that were stimulated for 60 seconds, cells in 100% calcium medium exhibited a greater response than both the 75% calcium and unstimulated control groups. Only ≈ 1% of cells responded under 75% calcium conditions, representing only a twofold increase over the unstimulated 100% calcium control (≈ 0.5%) (Figure 5K). Our results verify the role of calcium in the modulation of permeability. The effect of electrical stimulation on intestinal model permeability is dampened by the partial depletion of extracellular calcium, verifying that calcium entry into stimulated cells plays a major role in altering tight junction state, and therefore the paracellular permeability.

### In vivo validation of electrical stimulation model

We next sought to validate whether these effects are consistent in physiological tissue results in an acute in vivo set up. We used an acute in vivo model where anesthetized mice intestinal sections were exposed and 1 cm intestinal segments were placed in 24 well plate containing phosphate buffer solution. Sections were stimulated for 10 mins (1 mA, 500 Hz, 1 ms pulse width). FITC dextran tracers were injected into the intestinal lumen and the solution in each well sampled at different time points (Figure 6A). We measured fluorescence of the collected samples and compared the fluorescence to the transport kinetics of the control segment of the intestinal lumen. Concentration of FITC dextran rapidly increases after the first sampling of the solution 5 minutes post injection where it continues to increase upto ≈ 18.1 µM (Figure 6B). The apparent permeability of the electrically stimulated segment to FITC dextran is 8-fold larger compared to P_app_ = 0.9 µm/s compared to control (P_app_ = 0.9 µm/s vs 0.1 µm/s (Figure 6C)).

**Figure 6.**
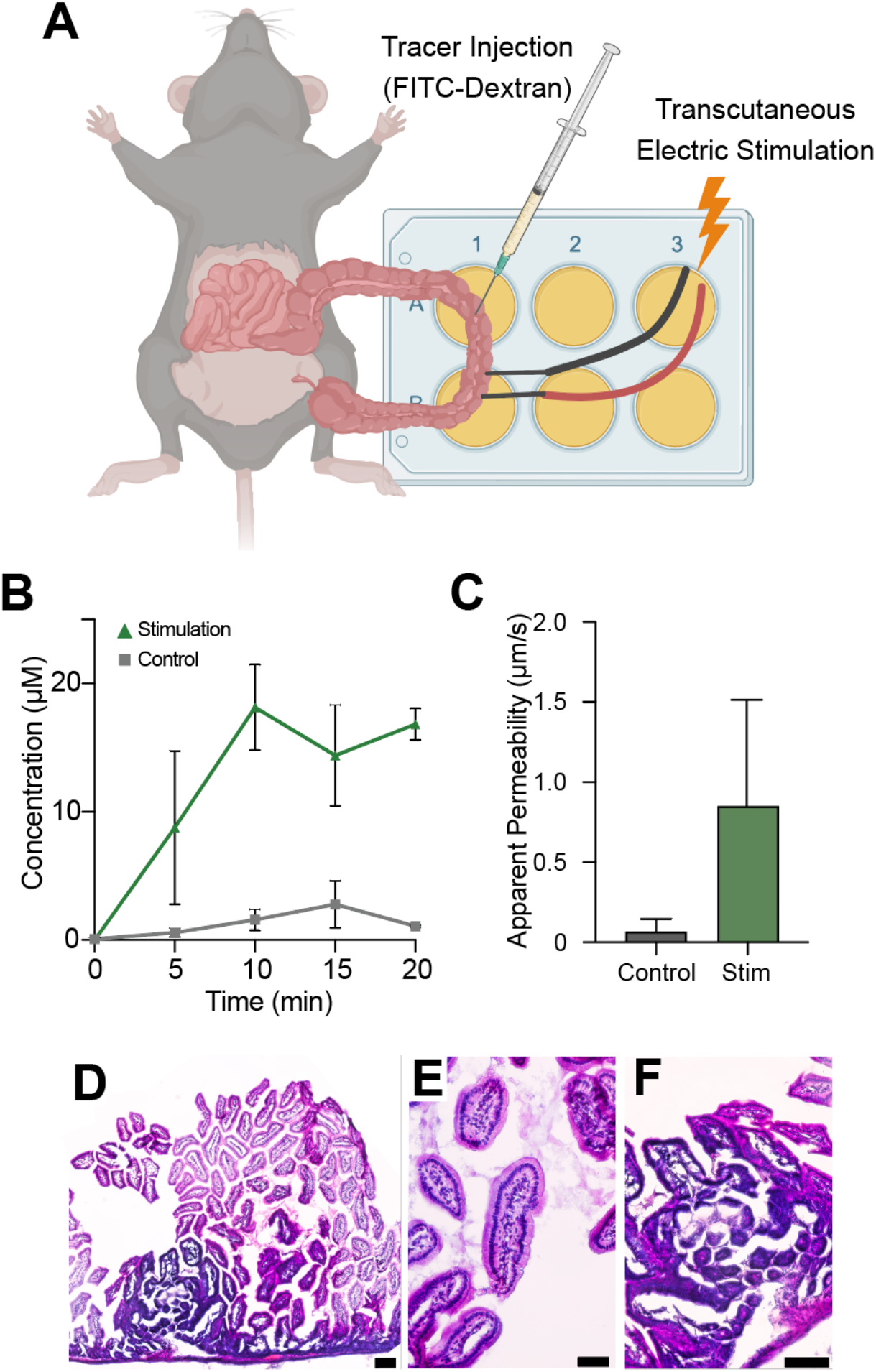
(A) Schematic of Ex-vivo Set up. (B) Concentration of FITC-Dextran in perfused intestinal epithelial samples in stimulated vs control sections, (C) Calculated apparent permeability of FITC Dextran in stimulated epithelial section vs control, (D) Hematoxylin and Eosin staining of sampled electrically stimulated large intestine (scale bar = 100 µm), (E) Colonic crypts of stimulated large intestine (Scale bar = 50 µm), (F) Basal crypts of stimulated large intestine (Scale bar = 50 µm).

Histological analysis of electrically stimulation tissue showed preservation of the overall intestinal architecture and intact epithelial morphology (Figure 6D). Stimulated regions did not exhibit signs of necrosis, cellular degeneration, or loss of tissue integrity (Figure 6E). Basal crypt structures were observed to be slightly distorted whilst maintaining their epithelial borders (Figure 6F). This suggests that the applied stimulation parameters did not induce significant pathological damage.

## DISCUSSION

We show how pulsed electrical stimulation can modulate gut epithelial barrier function. We quantify changes in permeability by measuring transepithelial electrical resistance (TEER), drug transport, as well as characterizing expression and organization of Zonula Occludens-1 (ZO-1), providing both functional and structural assessment of tight junction dynamics.

Studies on endogenous electrical fields that modulate cell functions have been previously reported, implicating the role of electric fields in cellular processes such as proliferation, migration, polarity and cytoskeletal organization[6, 14, 15]. Bioelectric cues play a role in modulating cell-cell interactions and junctional stability in epithelia, with previous studies demonstrating phenomena such as electro-inflation in MDCK organoids [16]. These studies support the concept that intestinal epithelial tissues are electrically responsive and that bioelectrical signals can influence tissue-level organization. Furthermore, reorganization of junctional complexes may be induced by membrane potential changes, which can alter the conformation of the proteins and associated cytoskeletal elements[17, 18]. Tight junction proteins, including ZO-1, interact with actin filaments in a charge- and tension-dependent manner, which suggests that electrical perturbation of endogenous fields may be implicated in the permeability modulation we have observed. This process is hypothesized to be related to calcium extracellular and intracellular signaling that directly influences cytoskeletal and tight junctional remodeling as studies have previously shown and our results corroborate[19-21]. While our findings demonstrate functional modulation of barrier integrity following pulsed electrical stimulation, further studies could directly measure changes in membrane potential and delineate their mechanistic contribution to tight junction remodeling.

The concept of using current to direct charged molecules across biological barriers has been previously applied in epithelial devices and commercially available cosmetic iontophoretic devices [13, 22]. These approaches rely on the iontophoretic principle, in which an electric field is applied across a barrier to electrophoretically drive charged molecules through tissue [23]. However, this model assumes controlled current directionality and uniform electric fields, which is not always practical in vivo. Our approach differs from these in kind, not merely in implementation. Rather than electrophoretically transporting a charged cargo while current flows, pulsed stimulation transiently and reversibly modulates the barrier itself. In our permeability assays the tracer is introduced only after stimulation has ended, so the increased transport we measure occurs with no field present and therefore cannot be attributed to iontophoretic drive. The effect is also cargo-agnostic: we observe enhanced transport of a large, near-neutral macromolecule (4 kDa FITC-dextran) that classical iontophoresis would not readily electromigrate. Because of this, we also show how barrier function can be modulated through conductive media and without electrodes coming into direct physical contact with epithelial cells.

While our study exhibits the effects of electrical stimulation on the increase of intestinal epithelial permeability, an increase in permeability may be indiscriminate and facilitate uptake harmful molecules. One way to limit uptake of undesired species could be colocalizing electrical stimulation to site of drug absorption. Further mapping the design space could also facilitate the identification of optimal parameters for localized, fast-reversible permeability increases. This may also reveal if there are unique stimulation parameters at which electrical fields interface with endogenous cell pathways to modulate tight junctions, to not only increase permeability but also decrease it. Our findings in the role of calcium in the downstream effects of electrical current on cellular pathways further emphasize the importance of investigating externally applied electrical stimuli effects on a systemic multicellular level. This could also provide a new perspective to disordered epithelial states where a permanent “leaky gut” is explored through the lens of a dysregulation in the membrane potential or endogenous electrical fields of the enterocytes.

## CONCLUSION

We demonstrate the potential of pulsed electrical current to modulate intestinal barrier function. By inducing controlled and reversible changes in permeability, this approach presents a promising strategy for applications including enhanced drug delivery. Beyond therapeutic delivery, electrically mediated barrier modulation may also serve as a novel research tool for studying bioelectrical cues in the gut, by controlling exogenous electric fields and examining its downstream effects on biological pathways involved in cell proliferation, regeneration and differentiation.

## METHODS

### Computer Simulations

The iontophoretic and direct stimulation models were set up on COMSOL Multiphysics 5.2. Material properties were set for the the electrodes and the polystyrene well material according to the material library. Media was set to have an electrical conductivity of 1.5 S/m and a relative permittivity of 60. The cell layer was set was have an electrical conductivity of 1×10^-5^ S/m and Relative permittivity of 500 to more closely simulate biophysical conditions. Next, the Electric Current module was used to set the boundary conditions (Terminal and Ground) of the electrodes and to allow for insulative conditions to be set for the well material. The bulk cell layer surface boundary was selected for a distributed impedance boundary condition and the surface resistance and capacitance were set to 1 Ω*m^2^ and 0.01 respectively. 1 ms pulsed current injected into the terminal was varied for the study and inputted as a waveform function.

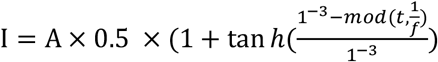 Where A is the current amplitude (amps),and *f* is the frequency (Hz)

A time-dependent study was then computed and the time was set to a range of (0, 2×10^-5^, 0.006) seconds for the 500 Hz and 1000 Hz and a range of (0,2×10^-5^, 0.15) and (0,2×10^-5^,0.02) seconds for 10 Hz and 100 Hz frequencies, respectively. Time stepping settings were set to have an initial step of 1×10^-6^ and a maximum step of 2×10^-5^ in order to mitigate non convergence of the solution. Voltage potential, electric field and current density norm results were probed at the bottom boundary of the cell layer and computed. Results were exported and plotted on PRISM 10.

### Cell Culture

Colorectal Adenocarcinoma cells (passages 3-32) were cultured in DMEM 10% Fetal Bovine Serum and 1% PenStrep. At 70% confluence, they were harvested and the cells were seeded on 24-well Transwell inserts at a density 1×10^5^/cm^2^ to differentiate for 21 days. Transepithelial Electrical Resistance (TEER) measurements were taken every 2 days and the media was changed every 2 days. Once the monolayers have reached a TEER equal to or above 400 Ω • cm^2^, they were used for further experiments.

### Electrical Stimulation Apparatus Construction

The apparatus was constructed by creating laser holes in 24-well plate lids that are 3 mm apart. Platinum Iridium 0.004” wires were then inserted into the holes, passed through pipette tips and were epoxied to the lid to ensure that the wires do not twist. The circuit was made to be a series circuit to allow for equal current distribution. The apparatus was then sprayed with 70% Ethanol and exposed under UV for 24 hours before being used to stimulate the Caco-2 monolayers.

### Electrical Stimulation Paradigms

Repeated measurements of TEER values were taken of the control and experimental wells of the Caco-2 monolayers before stimulation. The stimulation lid was placed on top of the experimental wells and the wires were confirmed to be submerged in the cell culture medium before being attached to positive and negative electrodes. Pulsed electrical stimulation was conducted using AM-Systems-2100 device. The parameters were varied either by time, frequency or amplitude. Following stimulation, repeated TEER measurements were taken of the control and stimulated wells and the resistance was calculated by multiplying the measured values by the growth area of the inserts (0.33 cm^2^). The change in TEER values before and after stimulation was then plotted for both control and stimulation groups.

For calcium depletion trials, Cells were incubated in a 1:1 (50%) and 3:4 (75%) Cell culture medium:HBSS-Ca^2+^-Free ratio for 20 minutes pre stimulation (N=2). TEER measurements were recorded following 20 minutes incubation. Cells were then stimulated for 10 minutes at a frequency of 500 Hz and 1 mA current amplitude. TEER measurements were recorded following stimulation and compared against the controls (N= 5) (HBSS-Ca^2+^-Free No stimulation and cell culture medium-Stimulation). One-way Brown-Forsythe ANOVA test was done on the ΔTEER values of the three experimental groups and Dunnet’s test was used to determine statistical significance using PRISM 10.

### Permeability Assay

Caco-2 monolayers were stimulated for 10 minutes at 1 mA, 500 Hz, 1 ms pulse width. The cell culture medium was then replaced with Hank’s Buffer Saline Solution (HBSS) in the basolateral layer. 250µL of 10 µM of Fluorescein was then prepared in HBSS and placed in the apical side. Following 20 minutes of incubation, 100 µL were sampled from the control and experimental wells’ basolateral side every 20 minutes for 120 minutes and 100 µL fresh HBSS was replaced in the basolateral layer at each timepoint. Standard curves were prepared and the samples’ fluorescence was read at excitation and emission wavelengths were read at 460 nm^2^ and 515 nm^2^ using the microplate reader (Biotek Cytation 5).

Apparent Permeability calculations were done by interpolating the unknown concentrations of the sample timepoints by plotting against the fluorescence values derived from the standard curve’s known concentration. Linear regression was then calculated and the derived slope was used to determine the Apparent permeability based on the below equation:

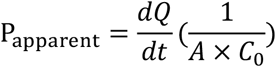

where 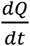 is the steady rate flow calculated from the slope multiplied by the receiver compartment’s volume, *A* is the area of well, and *C*_*0*_ is the initial concentration. Stimulated values were then normalized to the apparent permeability values of the control non stimulated cells, plotted on Prism 10, and compared across trials (N=2).

### Immunohistochemistry

Stimulated and control Caco-2 monolayers were gently washed with dPBS twice and fixed for 20 minutes using 4% Paraformaldehyde. Cell monolayers were then permeabilized using 0.2% Triton X for 10 minutes. Non-specific binding sites were blocked by incubating wells with 1% Normal Donkey Serum (NDS) in PBS for an hour. Zonula Occludin Primary antibody was prepared in 1% NDS at a ratio of 1:200 and the cells incubated in it overnight. Secondary antibody Alexa Fluor 594 (donkey anti-rabbit) was prepared in 1% NDS (1:100). The nucleus was stained using DAPI in PBS (1:1000). Following staining, the transwells were washed with PBS, cut out and placed on a glass slide and mounted using Fluorescence mounting medium. Acquired images were processed and analyzed using ImageJ. Localized thresholding was applied to all images and particle analysis was applied to the pores, counted and compared across controls and stimulated samples.

### Live Calcium imaging

Differentiated Caco-2 cells were loaded with 5 µM Oregon Green BAPTA-AM calcium dye and incubated for 60 minutes. Following that, de-esterification was carried out by incubating cells for 20 minutes in standard HBSS for both 100% and 75% calcium trials. To deplete calcium by 75% before imaging, Standard HBSS was switched with HBSS-Ca^2+^-Free medium prior to imaging.

Timelapse images of cells were captured using the Andor Dragonfly 600 Spinning Disk Confocal Microscope. 60 seconds control timelapse was taken prior to electrical stimulation exposure. Multiple regions were then recorded and exposed to stimulation duration at 1 mA, 500 Hz according to the stimulation paradigm in Table 2. Pre-stimulation recordings were used to normalize the data and post-stimulation recovery recordings were captured for 30 seconds post-stimulation.

**Table 2:**
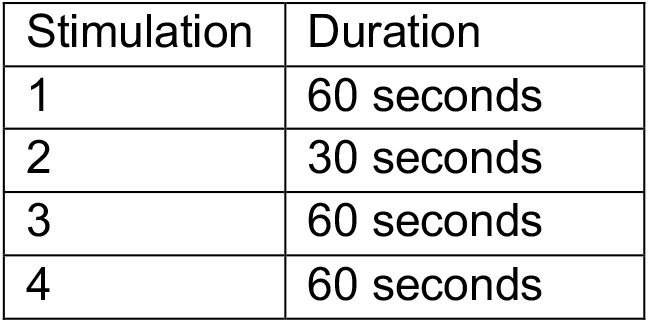
Stimulation paradigm of Caco-2 cells with multiple stimulation exposures and their durations.

| Stimulation | Duration |
| --- | --- |
| 1 | 60 seconds |
| 2 | 30 seconds |
| 3 | 60 seconds |
| 4 | 60 seconds |

Timelapse images of cells that were incubated in 75% HBSS-Ca^2+^ free medium were captured under the same acquisition settings and stimulated for 60 seconds at 1 mA, 500 Hz frequency. Pre-stimulation recordings were used to normalize the same and post-stimulation recovery recordings were captured for 30 seconds post stimulation.

### Image Processing of Calcium live images

Timelapse images were binned so that each frame = 1 second and imported into ImageJ as stacks. To segment the individual cells, a maximum projection Z-stack image was computed. On the projected image, an interactive watershed image was generated using the SCF-MP FIJI Plugin [24] (Supplementary Fig 4A). Following that, label size filtering was done (≥ 1500) on the generated image using the MorpholibJ Plugin. Morphological segmentation was then executed on the filtered image and an overlaid dams image was used to set threshold and segregate cells by their borders (Supplementary Fig4B-C)[25]. Particle analysis was then generated on the computed images and Rois were counted and the mean intensity was measured for all Rois (Supplementary Figure 4D-F)

### Data Processing of Calcium Live images

Mean fluorescence curves of cells fof the timelapse was measured and normalized according to the baseline fluorescence pre-stimulation.

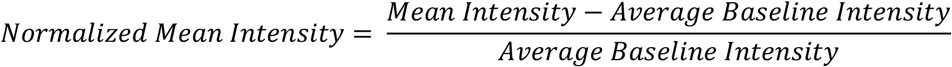

Signal to noise ratio (SNR) values were calculated for all timepoints and cell responses/peaks were filtered according to SNR ≥ 0.1. A secondary filter was run on the SNR-filtered data, where the maximum mean intensity ≥ 0.02.

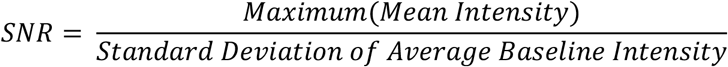

Response was characterized as the percentage of cells that produce peaks over the number of cells imaged in the field. Peaks of Calcium 100% and Calcium 75% filtered populations were then aligned according to the peak occurring at the earliest timepoint and plotted on Prism 10.

### Ex-vivo Stimulation

Adult mice were anesthetized and a midline laparotomy was performed to expose the small intestine. A 5–7 cm segment of the gut was exteriorized and gently laid flat into a sterile 4 well plate containing phosphate-buffered saline (PBS) to maintain hydration and to allow for collection of tracers. A 1 mA,500 Hz, 1ms pulsed stimulation was applied transcutaneously across a defined region of the intestine using sterile platinum/iridium electrodes for a total duration of 10 minutes. Adjacent, unstimulated regions of the intestine were used as internal controls. Following stimulation, 50µL FITC-dextran was injected into both the control and stimulated intestinal lumens. Aliquots (100 µL) of PBS from each were collected every 5 minutes over a 20-minute period to assess the appearance of tracer molecules in the surrounding solution, indicating enhanced permeability. Fluorescence intensity was measured on a microplate reader, and concentrations were determined based on standard curves. Intestinal regions were harvested and immersed in 4% Paraformaldehyde Evaluation of stimulated Intestinal regions was done by embedding fixed samples in cryo-embedding medium (Tissue-Tek) prior to cryosectioning. Longitudinal sections of ≈ 15 µm were captured using a Leica CM1950 Cryostat. Staining for Hematoxylin and Eosin was done by dehydrating samples and incubating them in Hematoxylin and Eosin according to manufacturer protocol (AB245880). Slides were imaged using Nikton Ti-2 Widefield Microscope (Nikon Instruments Inc.) and Z stack images were processed using FIJI.

## ACKNOWLEDGEMENT

This work was supported by the Center for Translational Medical Devices and Center for Brain Health at NYU Abu Dhabi. We acknowledge NYU Abu Dhabi Core Technology Platforms for equipment and Dr. Rachid Rezgui’s technical assistance.

## DATA AVAILABILITY STATEMENT

All data that support the findings of this study are included within the article (and any supplementary files).

## ETHICS STATEMENT

All animal experiments were performed in accordance with protocols approved by the Institutional Animal Care and Use Committee at New York University Abu Dhabi (Protocol 22-0006).

---

**Figure S1.**
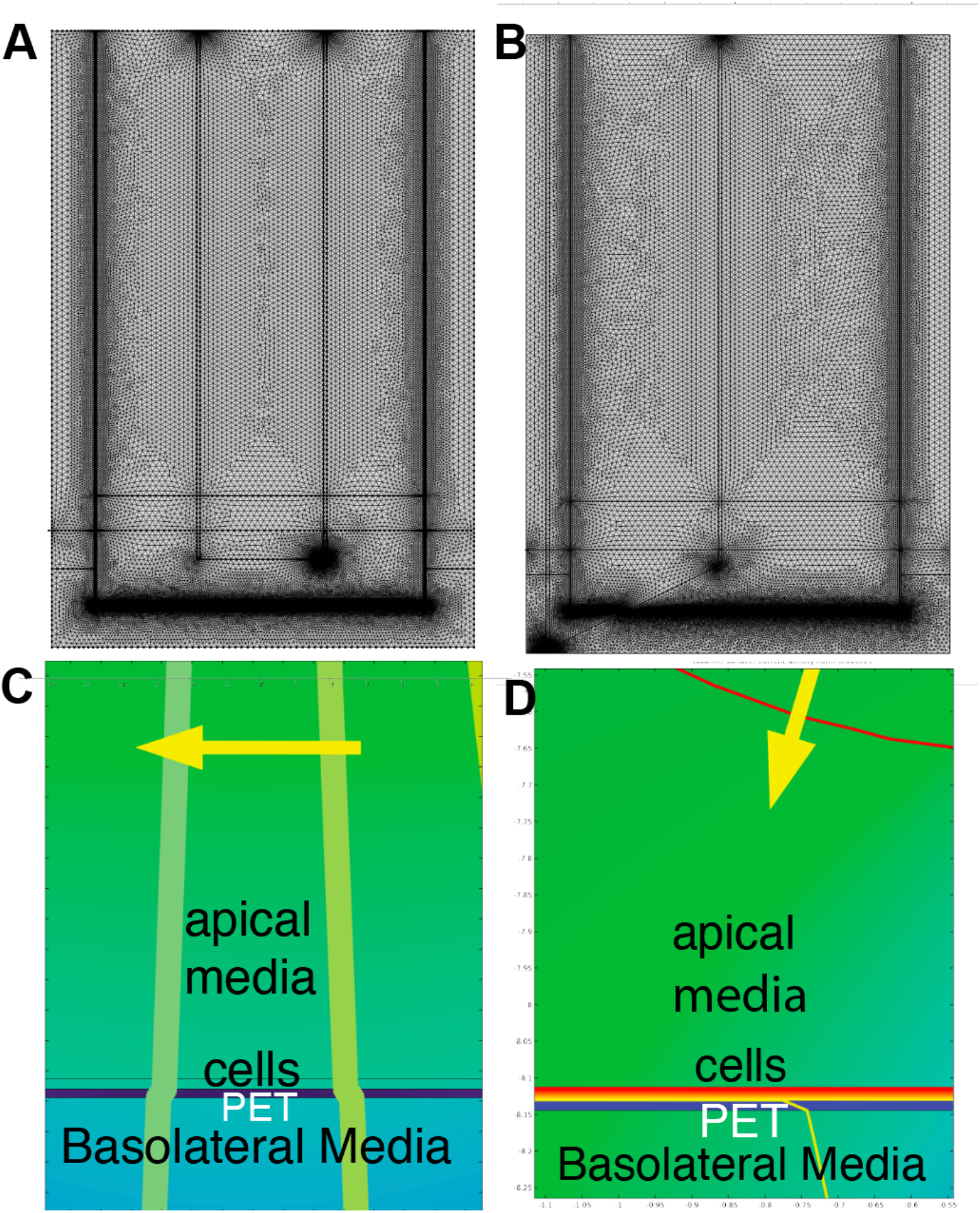
(A-B) Mesh of (A) direct stimulation and (B) iontophoretic COMSOL simulation. (C-D) Close up of the cell layer during (C) direct and (D) iontophoretic Simulation

**Figure S2.**
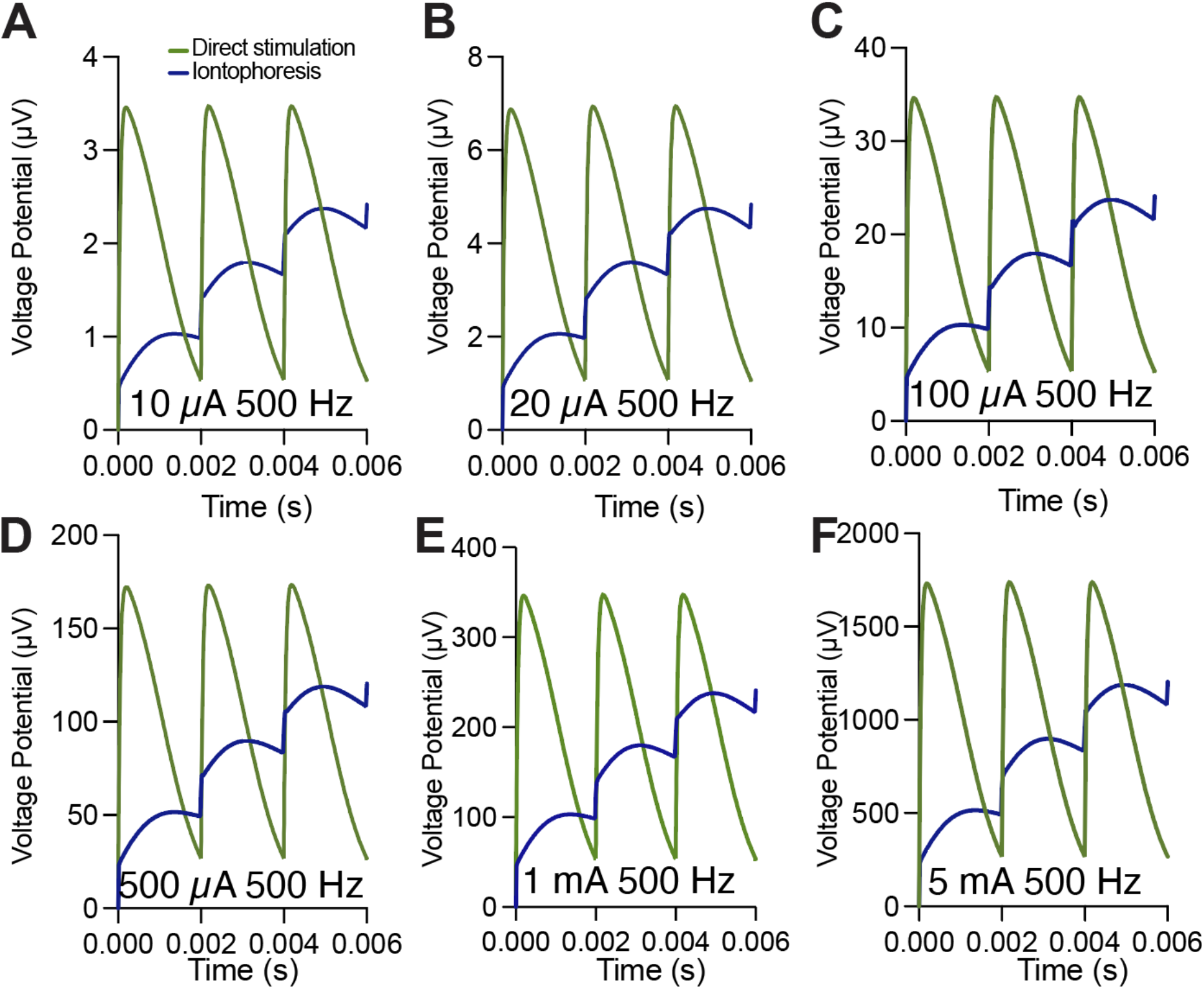
(A) Voltage potential of cell layer at 10 µA 500 Hz stimulation, (B) Voltage potential of cell layer at 20 µA 500 Hz stimulation, (C) Voltage potential of cell layer at 100 µA 500 Hz stimulation, (D) Voltage potential of cell layer at 500 µA 500 Hz stimulation, (E) Voltage potential of cell layer at 1 mA 500 Hz stimulation, (F) Voltage potential of cell layer at 5 mA 500 stimulation

**Figure S3.**
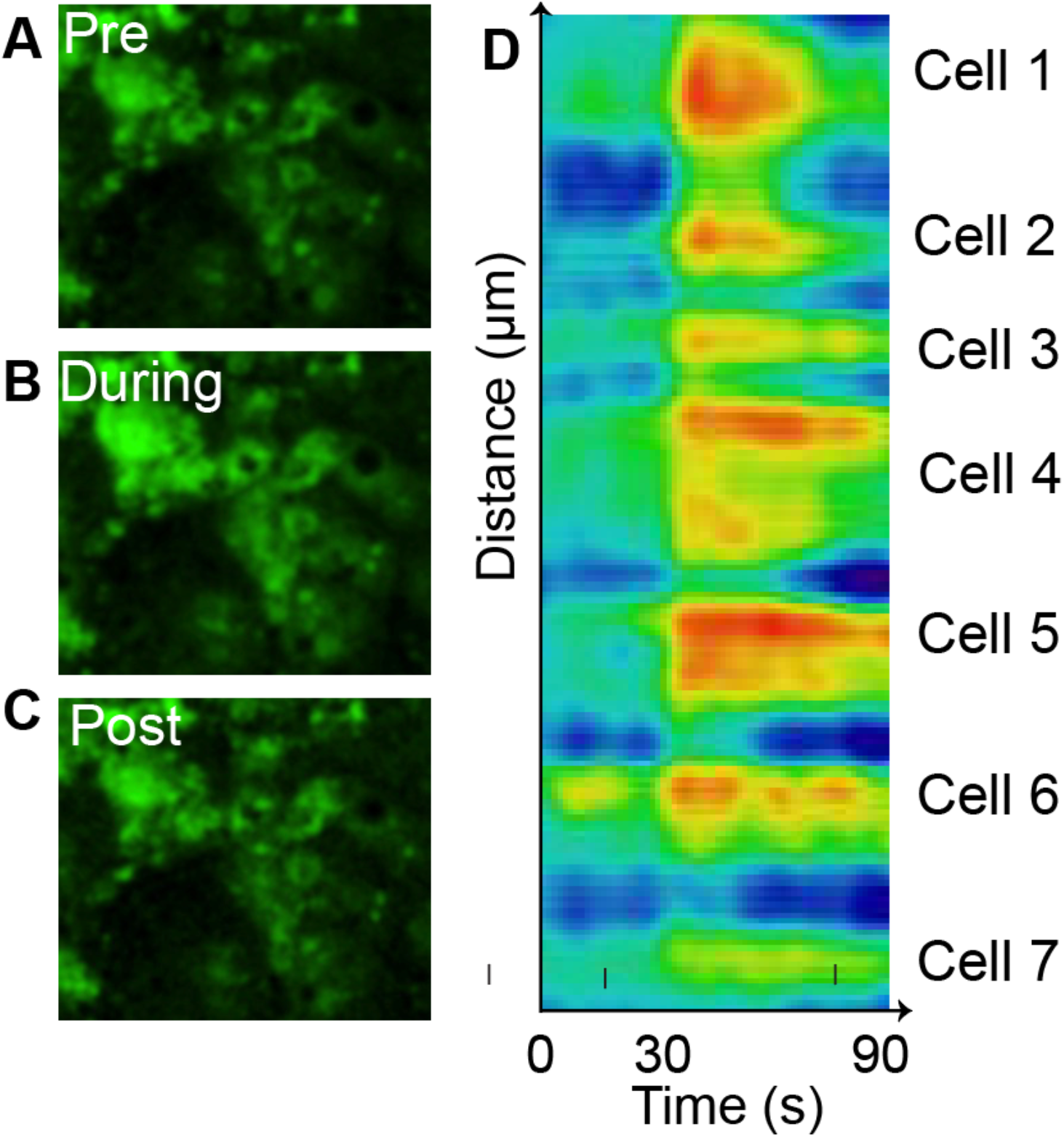
(A) Calcium image of a population of cells pre-stimulation, (B) Calcium image of a population of cells during stimulation, (C) Calcium image of a population of cells post-stimulation, (D) Kymograph of the population of cells showing spontaneous signaling of cells.

**Figure S4.**
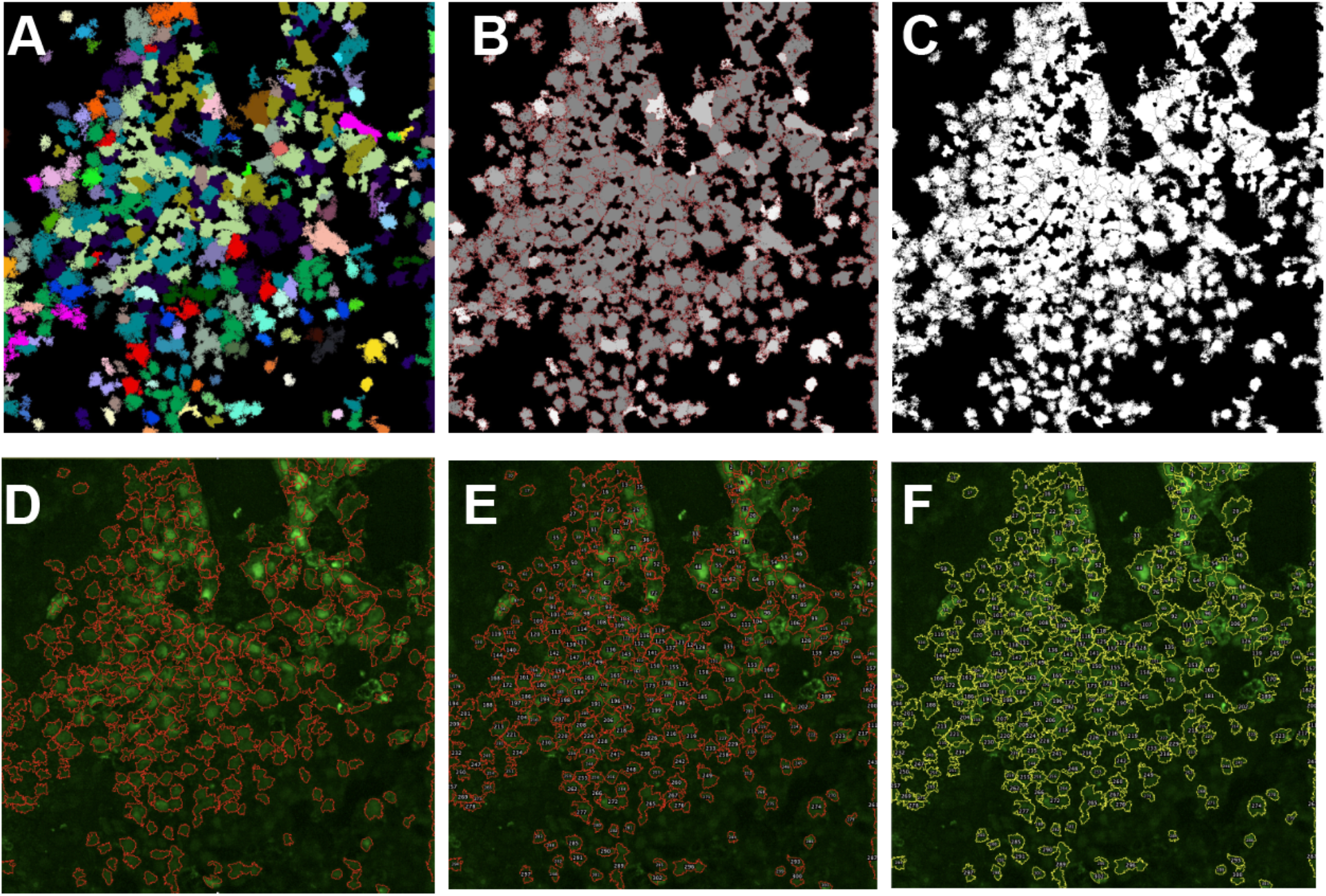
(A) Interactive watershed image generated, (B) MorphoLibJ Morphological segmentation, (C) Binary Image of segmented cells, (D) Particle Analysis of Segmented cells, (E) Count of segmented cells, (F) ROI labels over segmented cells

